# Comparative genomics and structure-informed analysis identify a putative tandem receptor-binding module in *Pseudomonas* phage UF_RH1

**DOI:** 10.64898/2026.10.07.757288

**Authors:** Golnaz Ebrahimi, Mehdi Mirsaeidi

**Affiliations:** R&D Division, Phageolytix Inc., Coral Springs, Florida, USA; Division of Pulmonary and Critical Care Medicine, Dell Medical School, The University of Texas at Austin, Austin, Texas, USA

**Author notes:** Corresponding author: Golnaz Ebrahimi, MD.

**Keywords:** bacteriophage, Pseudomonas aeruginosa, UF_RH1, Septimatrevirus, receptor-binding protein, tail spike, type IV pilus, phage genomics, host range

## Abstract

Bacteriophage receptor-binding proteins (RBPs) determine host recognition and host range but are frequently missed by conventional annotation because they evolve rapidly and often lack recognizable sequence domains. Pseudomonas phage UF_RH1 is a lytic Septimatrevirus with a 42,567-bp double-stranded DNA genome encoding 57 predicted open reading frames, several of which remain annotated as hypothetical proteins. Here, we reanalyzed the UF_RH1 distal-tail region using comparative synteny, phage RBP-specific prediction tools, structure-informed domain analysis, oligomeric-symmetry prediction, and independent physicochemical calculations. Two adjacent proteins, WCF59042.1 (317 aa) and WCF59043.1 (320 aa), emerged as the strongest candidates for functional reannotation. WCF59042.1 was classified by DepoScope as a tail-spike protein with probability 1.00 and contains a high-confidence C-terminal tail-spike-associated segment spanning residues 210–317. WCF59043.1 was independently classified as a tail-spike candidate by RBPdetect2 (probability 0.51) and showed a strong Seq2Symm prediction for C3 homo-trimeric symmetry (0.990). Comparative genomic analysis places these proteins at positions corresponding to DLP2 gp22 and gp23, two similarly sized hypothetical virion proteins immediately downstream of the tape-measure gene in a conserved distal-tail morphogenesis cassette. Because the closely related phages DLP1 and DLP2 use type IV pili for adsorption, we propose a testable model in which WCF59042.1 functions as a proximal connector/adaptor and WCF59043.1 contributes a distal trimeric receptor-binding element. The model is computational and does not establish direct binding to PilA or assembled pili. Together, the convergent sequence, structural, oligomeric, and syntenic evidence supports a previously unrecognized tandem distal-tail receptor-binding module in UF_RH1 and provides experimentally tractable targets for defining Septimatrevirus adsorption and host range.

## Introduction

Bacteriophages initiate infection by recognizing specific structures exposed at the bacterial surface through virion-associated receptor-binding proteins (RBPs).^1,2^ These receptors include lipopolysaccharides, capsular polysaccharides, membrane proteins, pili, and flagella.^1–3^ Because adsorption is the first committed step of productive infection, variation in RBPs can profoundly alter host specificity even among otherwise closely related phages.^4,5^

RBP annotation remains difficult. Receptor-binding proteins often show substantial primary-sequence divergence, reflecting rapid adaptation to variable bacterial surface receptors and host range.^4,6^ Tail fibers and tail spikes also exhibit modular, repetitive, and structurally diverse architectures that can be poorly represented in conventional sequence-domain databases.^6^ As a result, functionally important virion proteins may remain designated simply as hypothetical proteins after complete genome sequencing.

We previously reported the complete genome of the lytic Pseudomonas aeruginosa phage UF_RH1.^7^ UF_RH1 contains a linear, circularly permuted 42,567-bp double-stranded DNA genome with 53.56% GC content and 57 predicted open reading frames. Comparative analysis placed UF_RH1 within the genus Septimatrevirus and identified Stenotrophomonas phage vB_SmaS-DLP_2 (DLP2) as a close relative, with 97% genome coverage and 97.8% nucleotide identity. The original UF_RH1 annotation identified proteins involved in genome replication, DNA packaging, virion morphogenesis, and host lysis, including DNA polymerase, DEAD-box helicase, RecB exonuclease, tape-measure protein, central tail hub, holin, and endolysin. However, multiple proteins in the structural region remained functionally unknown.

DLP2 and the closely related DLP1 provide a particularly informative comparator. Their genomes contain a conserved distal-tail cassette, and experimental studies demonstrated that both phages depend on type IV pili for productive adsorption.^8,9^ Deletion of the major pilin gene pilA abolished binding and infection, while complementation restored susceptibility.^10^ Despite this receptor-level information, the specific viral proteins mediating pilus recognition have not been firmly assigned.

We therefore revisited the UF_RH1 distal-tail region using contemporary phage-specific prediction resources and comparative genomic context. The central objective was not simply to relabel hypothetical proteins, but to identify a coherent structural module supported by independent evidence. Our results converge on two adjacent proteins, WCF59042.1 and WCF59043.1, and support a mechanistic model in which they form a tandem distal-tail connector/receptor-binding apparatus.

## Materials and Methods

### Genome and protein sequence sources

The complete UF_RH1 genome and protein annotations were obtained from GenBank accession OQ259603.1. The primary proteins analyzed were WCF59042.1 and WCF59043.1. WCF59052.1/UFRH1_26 was retained as a secondary unresolved hypothetical protein for comparison but was not used to support the receptor-binding model.

### Comparative genomic and synteny analysis

UF_RH1 genomic organization was compared with related Septimatrevirus phages, with particular emphasis on DLP2 (GenBank KR537871), whose genome architecture and predicted gene products have been characterized in detail.^8^ Genomic position, coding orientation, protein length, neighboring genes, and historical annotation were considered jointly. Because UF_RH1 is circularly permuted, gene number alone was not treated as sufficient evidence of orthology. Particular attention was given to the region immediately downstream of the tape-measure protein and upstream of the central tail-hub gene.

### RBP and domain prediction

UF_RH1 proteins were queried in the Phage Receptor-Binding Protein Database (PhaRBP; accessed October 3, 2026). Tail-spike/depolymerase predictions were evaluated using DepoScope, a protein-language-model-based approach for phage depolymerase classification and domain delineation.^11^ RBPdetect2 classifications available through PhaRBP were recorded for WCF59043.1. For each candidate, the predicted RBP class, model probability, sequence, available structural prediction, domain segmentation, and database metadata were reviewed.

### Structural and oligomeric-state information

Structure models available through PhaRBP were generated using ESMFold or ESMFold2. For WCF59042.1, the reported tail-spike domain segmentation was reviewed, including the high-confidence C-terminal segment spanning residues 210–317. For WCF59043.1, the reported ESMFold2 mean pLDDT and Seq2Symm probabilities were recorded. Seq2Symm predicts likely homo-oligomeric symmetry directly from protein sequence using protein foundation-model representations.^12^ The resulting symmetry probabilities were interpreted as architectural evidence rather than direct proof of biological oligomerization.

### Physicochemical analysis

Protein sequence properties were independently calculated using Biopython version 1.86 and Bio.SeqUtils.ProtParam.^13,14^ Calculated parameters included sequence length, molecular mass, predicted isoelectric point, aromaticity, GRAVY hydropathy, instability index, amino-acid composition, and selected acidic/basic residue counts. Exact input sequences, outputs, analysis dates, and a reproduction script are provided in Supplementary Table S1 and the accompanying supplementary data package.

### Mechanistic modeling and microscopy context

Mechanistic figures were constructed as hypothesis-generating schematics that integrate the computational reannotation with previously established type IV pilus receptor biology in DLP1/DLP2.^9^ These figures do not represent experimentally determined UF_RH1 protein–pilus contacts, a solved multimolecular complex, or experimentally measured kinetics. Downstream stages of DNA injection, intracellular replication, virion assembly, and lysis are presented as a canonical lytic-cycle framework rather than as experimentally mapped UF_RH1-specific intermediates.

### Transmission electron microscopy

Transmission electron microscopy was performed at the Indiana Center for Electron Microscopy (iCEM) using a Tecnai G2 Spirit BioTWIN transmission electron microscope. Purified bacteriophage preparations were applied to 300-mesh nickel Formvar/carbon grids, glow discharged for 25 s at 25 mA, and negatively stained with 2% aqueous uranyl acetate. Thin sections of phage-exposed bacteria were mounted on 200-mesh hexagonal copper grids and stained with 2% uranyl acetate in 50% ethanol. Images were acquired over a range of magnifications to evaluate phage morphology, bacterial ultrastructure, and phage–cell interactions. The original report noted fixation-associated aggregation and structural distortion in some phage preparations.

## Results

### The UF_RH1 distal-tail region contains a conserved cassette of unresolved virion proteins

UF_RH1 contains 57 predicted open reading frames spanning lysis, DNA packaging, capsid and tail morphogenesis, and replication/repair functions. Despite extensive genome-level conservation with DLP2, several proteins in the tail region remain annotated as hypothetical. DLP2 provides an informative reference because gp21–gp26 form a conserved distal-tail cassette: the 1,187-aa tape-measure protein gp21 is followed by gp22 (320 aa), gp23 (321 aa), gp24 (570 aa), and gp25 (274 aa), all historically annotated as hypothetical proteins with virion-protein similarity, and then by the 813-aa central tail-hub protein gp26.^8^

WCF59042.1 and WCF59043.1 occupy the corresponding UF_RH1 interval and closely match the sizes of DLP2 gp22 and gp23. Their tandem genomic placement immediately downstream of the tape-measure gene provides a strong positional argument that they are components of a conserved distal-tail assembly rather than unrelated accessory proteins (Figure 1).

**Figure 1.**
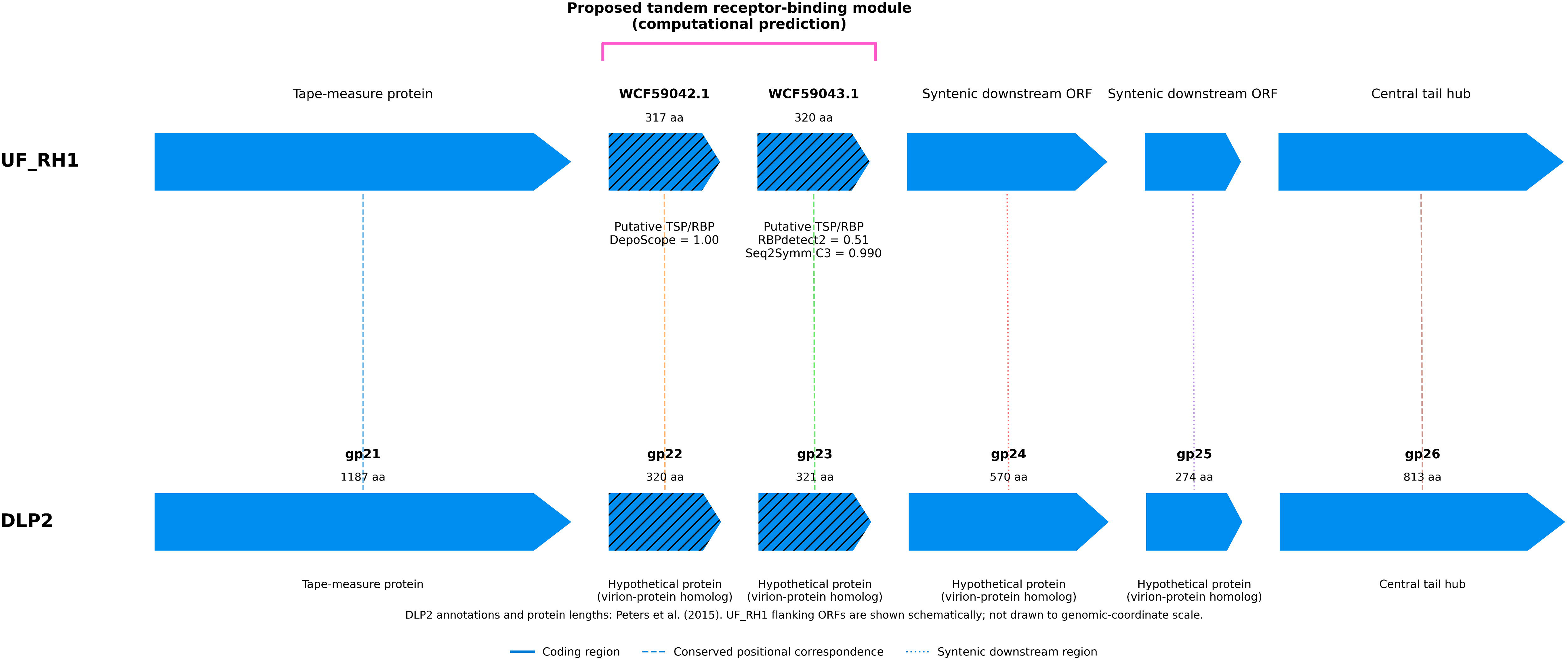
Conserved distal-tail synteny between UF_RH1 and DLP2. WCF59042.1 and WCF59043.1 occupy positions corresponding to DLP2 gp22 and gp23 within the distal-tail morphogenesis cassette immediately downstream of the tape-measure protein. The two UF_RH1 proteins are highlighted as the proposed tandem receptor-binding module. DLP2 protein lengths and historical annotations are from Peters et al.^8^ UF_RH1 flanking genes are shown schematically; arrow widths are not genomic-coordinate scale.

### WCF59042.1 is a high-confidence tail-spike candidate with a defined C-terminal segment

WCF59042.1 is a 317-aa hypothetical protein encoded at nucleotides 16,471–17,424 on the reverse strand. Independent ProtParam analysis yielded a molecular mass of 33.50 kDa, predicted pI of 8.19, and GRAVY value of -0.073. PhaRBP classifies WCF59042.1 as a tail-spike protein on the basis of DepoScope with probability 1.00. Conventional InterPro annotation is absent, highlighting the limitation of standard domain-based approaches for this protein.

Tail-spike segmentation divides WCF59042.1 into an N-terminal segment (residues 1–10), a central segment (11–209), and a C-terminal segment (210–317). The C-terminal segment carries the strongest segmentation confidence (0.9995), whereas the first two boundaries are considerably less confident. Accordingly, the most defensible structure-informed conclusion is that residues 210–317 constitute a strongly supported tail-spike-associated region rather than that the complete domain architecture is experimentally established (Figure 2).

**Figure 2.**
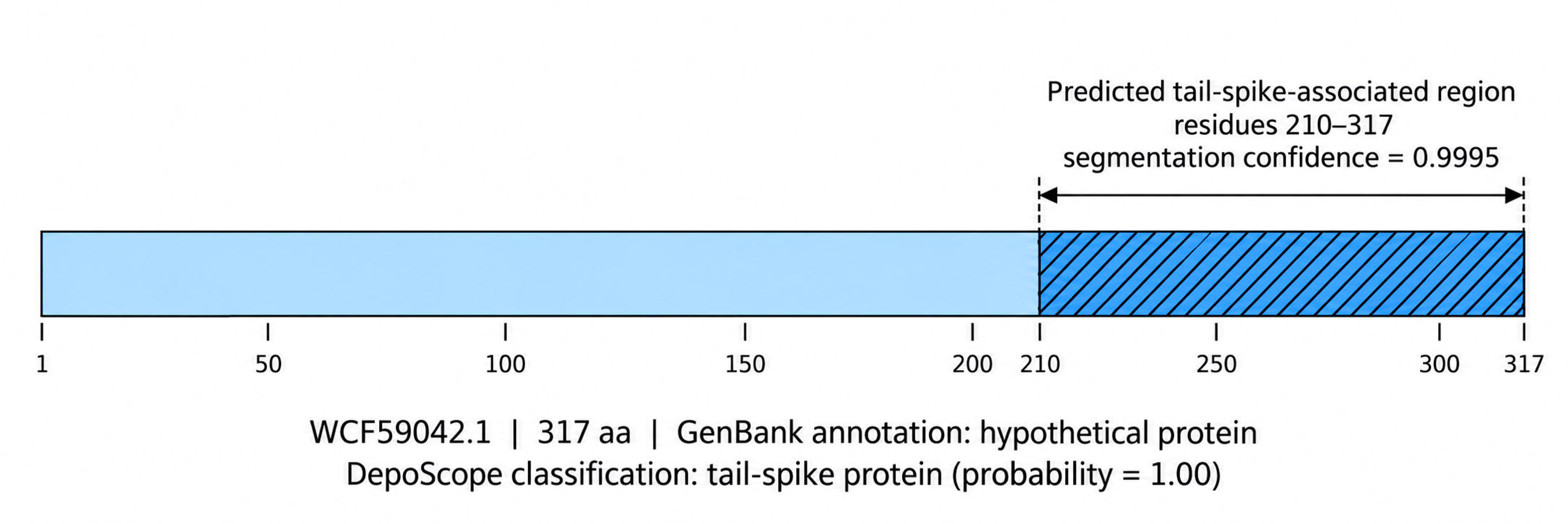
Structure-informed annotation of WCF59042.1. Sequence-level representation of the 317-aa protein showing the strongly supported C-terminal tail-spike-associated segment spanning residues 210–317 (segmentation confidence 0.9995). DepoScope classifies WCF59042.1 as a tail-spike protein with probability 1.00. The figure is a domain schematic and does not depict an experimentally determined structure.

### WCF59043.1 is a second RBP candidate with strong predicted C3 symmetry

WCF59043.1 lies immediately adjacent to WCF59042.1 and encodes a 320-aa hypothetical protein. Its calculated molecular mass is 34.99 kDa, predicted pI 5.24, and GRAVY value -0.033. RBPdetect2 classifies WCF59043.1 as a tail-spike candidate with probability 0.51. The moderate classification score alone would be insufficient for strong functional assignment.

However, two independent features strengthen the interpretation. First, an ESMFold2 model is available with mean pLDDT 73.2, consistent with a substantially ordered protein. Second, Seq2Symm predicts C3 homo-trimeric symmetry with probability 0.990, compared with 0.014 for C1 and 0.002 for C2 (Figure 3). Because trimeric architecture is common among phage tail-spike and receptor-binding proteins, the C3 prediction provides orthogonal support for a distal receptor-binding role, although it does not establish the actual oligomeric state in the virion.

**Figure 3.**
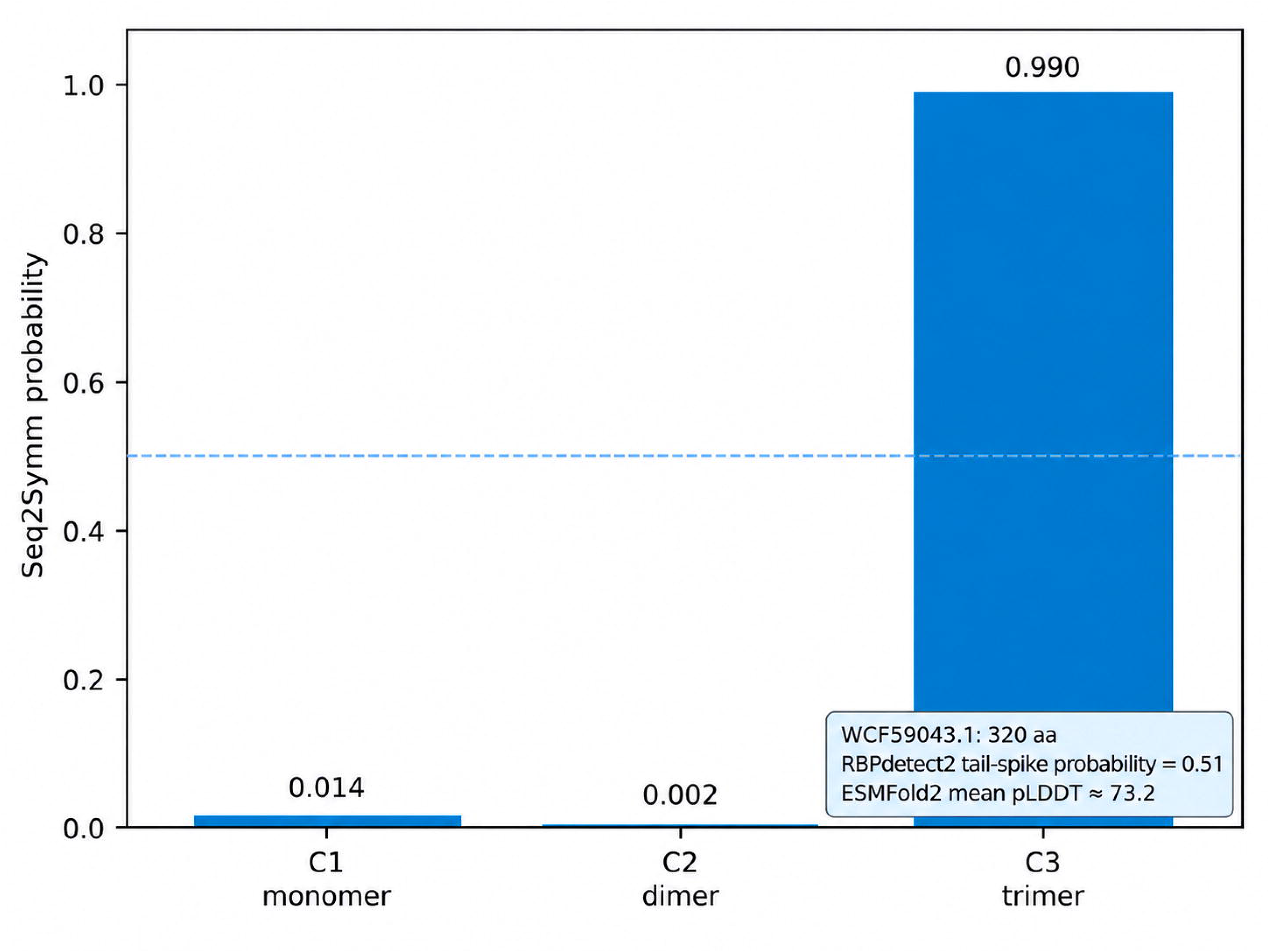
Predicted oligomeric symmetry of WCF59043.1. Seq2Symm assigns a C3 homo-trimeric symmetry probability of 0.990, compared with 0.014 for C1 and 0.002 for C2. WCF59043.1 is a 320-aa hypothetical protein classified by RBPdetect2 as a tail-spike candidate with probability 0.51; its ESMFold2 mean pLDDT is 73.2.

### Convergent evidence supports a tandem receptor-binding/connector module

The strongest interpretation emerges from combining gene order, protein size, historical virion association, RBP-specific predictions, and predicted oligomerization. WCF59042.1 and WCF59043.1 are adjacent, are positioned directly downstream of the tape-measure protein, correspond in size and position to DLP2 gp22/gp23, and are independently identified as tail-spike candidates by different computational methods.

These observations favor a model in which the two proteins perform complementary rather than redundant functions. We propose that WCF59042.1 may act as a proximal adaptor or connector between the conserved tail apparatus and a distal receptor-binding element, whereas WCF59043.1 may contribute a trimeric distal recognition component. This division of labor remains a working model; no docking experiment, cryo-EM reconstruction, or biochemical interaction study currently establishes direct contact between the two proteins.

### A type IV pilus–centered adsorption model is experimentally testable

DLP1 and DLP2 require type IV pili for adsorption, and their close evolutionary relationship to UF_RH1 makes pilus recognition a biologically plausible hypothesis for UF_RH1.^8,9^ We therefore propose a staged adsorption model in which the distal WCF59042.1/WCF59043.1 module encounters an exposed type IV pilus, WCF59043.1 contributes the distal recognition interface, and WCF59042.1 stabilizes or couples this interaction to the phage tail. Pilus retraction could then move the adsorbed virion toward the cell envelope, permitting irreversible engagement and genome delivery (Figure 4).

**Figure 4.**
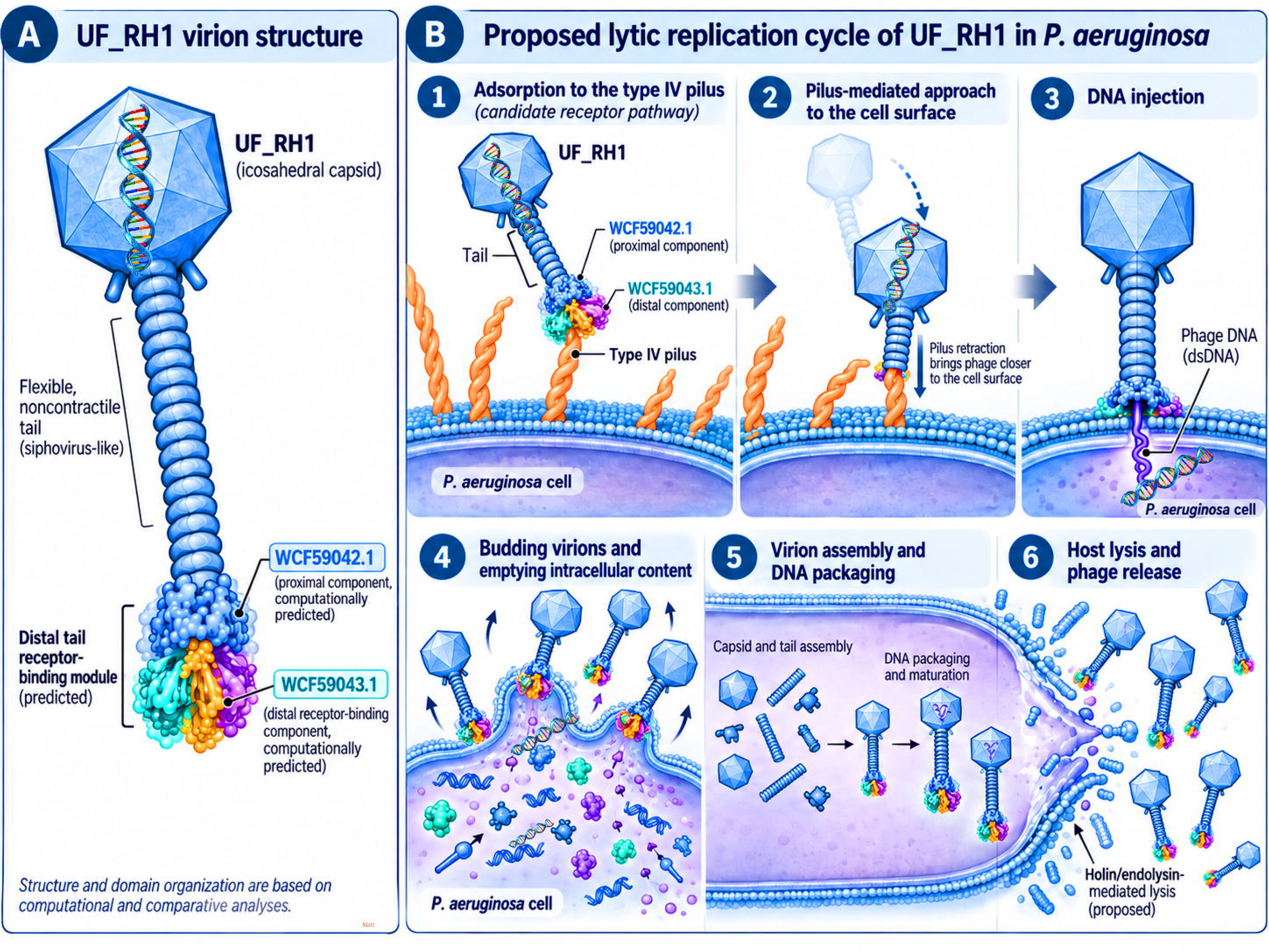
Proposed mechanism of UF_RH1 interaction with the P. aeruginosa type IV pilus. Working model integrating the predicted tandem distal-tail module with the experimentally established type-IV-pilus dependence of related DLP1/DLP2 phages. WCF59043.1 is depicted as the distal recognition component and WCF59042.1 as a proximal connector/adaptor. Pilus retraction is proposed to bring the adsorbed phage toward the cell surface. None of these UF_RH1-specific protein–pilus interactions has yet been experimentally demonstrated. Schematic figure design, conceptual layout, and selected graphical elements were developed with assistance from generative AI and subsequently refined, labeled, and assembled in Adobe Photoshop. All scientific content and final figure interpretation were reviewed and approved by the authors.

Only the comparative and computational components of this model are demonstrated here. UF_RH1 binding to PilA, pilus dependence, the directionality of WCF59042.1/WCF59043.1 within the mature virion, and the role of pilus retraction remain hypotheses requiring experimental validation.

### The predicted receptor-binding module can be placed within a broader lytic-cycle framework

To place the adsorption hypothesis in biological context, we developed a schematic lytic-cycle model (Figure 5). The figure separates the central mechanistic hypothesis, pilus engagement and pilus-mediated approach, from downstream canonical lytic events: genome injection, early gene expression and genome replication, virion assembly and DNA packaging, and holin/endolysin-mediated lysis with phage release. These later stages are consistent with the UF_RH1 genomic repertoire but are not individually resolved by the current computational RBP analysis.

**Figure 5.**
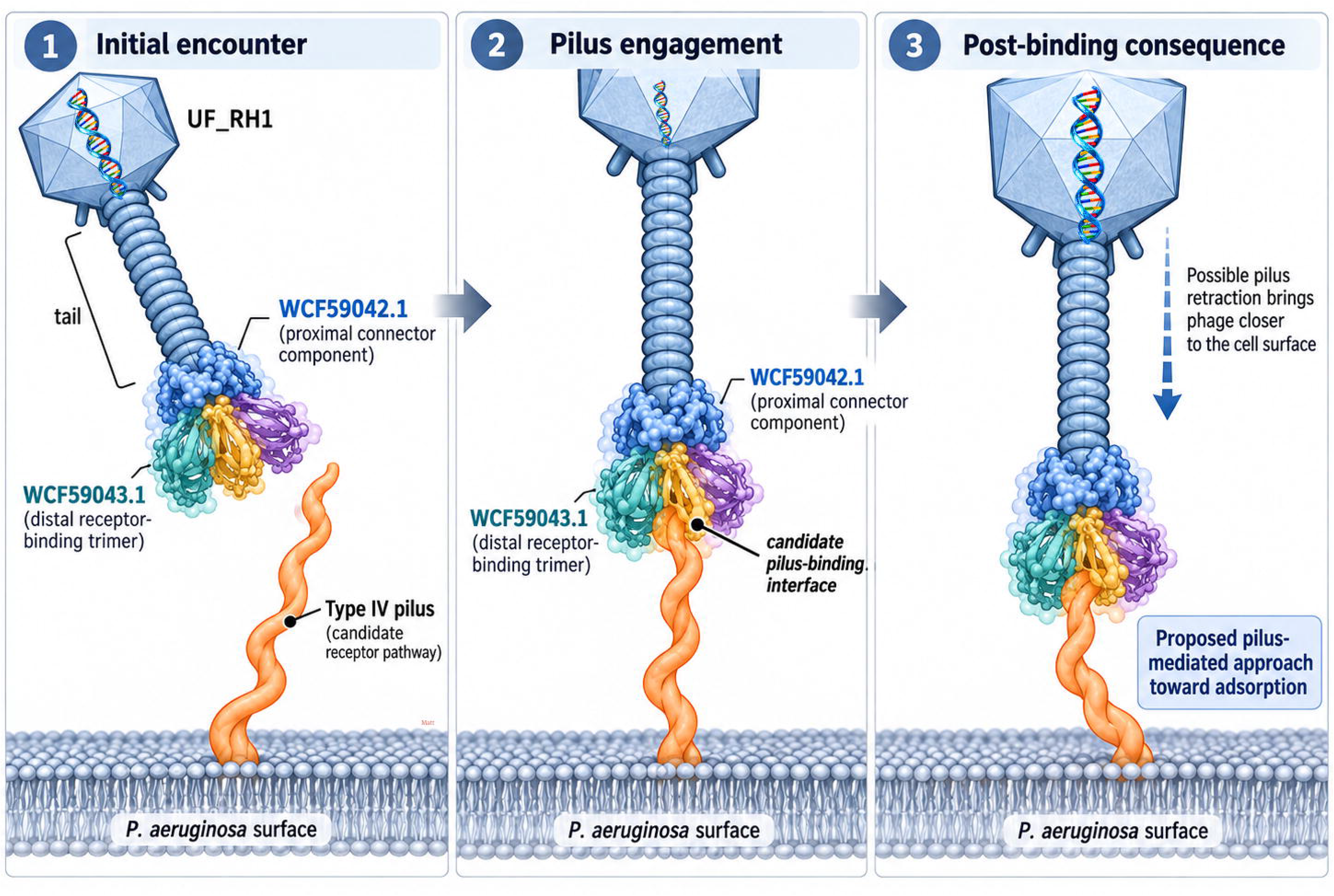
Proposed lytic replication cycle of UF_RH1 in P. aeruginosa. Schematic placing the receptor-binding hypothesis within a canonical lytic infection sequence: adsorption to a candidate type IV pilus, pilus-mediated approach to the cell envelope, DNA injection, early gene expression and genome replication, virion assembly and DNA packaging, and host lysis with progeny release. The downstream intracellular steps are biological context rather than direct outputs of the current RBP analysis. Schematic figure design, conceptual layout, and selected graphical elements were developed with assistance from generative AI and subsequently refined, labeled, and assembled in Adobe Photoshop. All scientific content and final figure interpretation were reviewed and approved by the authors.

**Figure 6.**
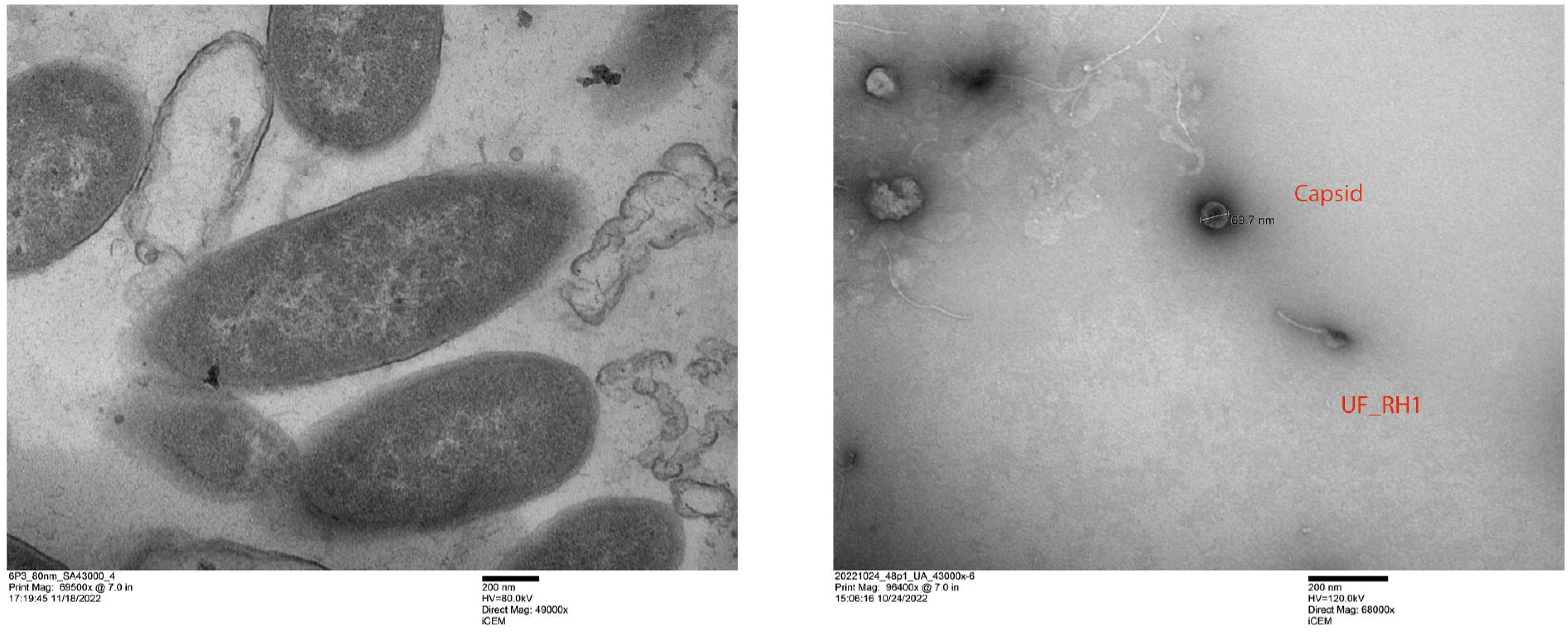
Transmission electron microscopy of UF_RH1 and UF_RH1-infected Pseudomonas aeruginosa. The left panel shows thin-section TEM images of infected P. aeruginosa representing different stages of phage-associated cellular disruption, ranging from active progeny-phage release, through partial loss of cell-wall integrity, to advanced collapse of the bacterial envelope. The right panel shows the morphology of purified UF_RH1, demonstrating a distinct capsid and associated tail; the measured capsid diameter is approximately 69.7 nm.

### UFRH1_26 remains unresolved and should not be conflated with the receptor-binding module

WCF59052.1/UFRH1_26 is a 131-aa acidic hypothetical protein with a calculated molecular mass of 13.45 kDa and predicted pI of approximately 4.05. It contains 21 Asp/Glu residues and only five Lys/Arg/His residues. Unlike WCF59042.1 and WCF59043.1, it lacks convergent RBP, synteny, or oligomeric evidence supporting a distal-tail assignment. We therefore retain it as a small acidic protein of unresolved function and exclude it from the principal receptor-binding model.

## Discussion

The original description of UF_RH1 established this isolate as a lytic *Pseudomonas aeruginosa* bacteriophage with a 42,567-bp double-stranded DNA genome and placed it within the genus *Septimatrevirus*.^7^ That initial genomic study successfully identified major genes involved in replication, morphogenesis, and host lysis, but several proteins within the structural region remained annotated as hypothetical. The present study extends the original genomic characterization by applying contemporary receptor-binding protein prediction, structure-informed analysis, oligomeric-state prediction, and comparative synteny to this unresolved portion of the UF_RH1 genome.

The principal finding is the convergence of several independent observations on two adjacent proteins, WCF59042.1 and WCF59043.1. WCF59042.1 received a high-confidence DepoScope tail-spike classification and contains a strongly supported C-terminal tail-spike-associated region, whereas WCF59043.1 was independently identified as an RBP candidate and showed a strong prediction for C3 homo-trimeric symmetry. This combination is biologically plausible because phage receptor-binding proteins are frequently located at the distal tail and commonly form highly organized multimeric assemblies that mediate host recognition and irreversible adsorption.^15,16^ Recent structural studies have further demonstrated that receptor engagement can induce substantial conformational changes within distal-tail complexes that couple recognition of the bacterial receptor to subsequent genome delivery.^16^

Importantly, these computational results are supported by conserved genomic organization. WCF59042.1 and WCF59043.1 occupy positions corresponding to the similarly sized DLP2 proteins gp22 and gp23 immediately downstream of the tape-measure gene. In the original DLP2 genome study, gp22 and gp23 were classified broadly as hypothetical or virion-associated proteins rather than being assigned specific receptor-binding functions. The correspondence in protein size, tandem organization, and location within the distal-tail morphogenesis region therefore provides an independent genomic framework supporting functional reannotation of the UF_RH1 proteins.

A tandem receptor-binding complex is more likely than two redundant tail spikes

Although WCF59042.1 and WCF59043.1 are similar in length, they differ substantially in primary sequence and physicochemical characteristics. WCF59042.1 is relatively basic, whereas WCF59043.1 is mildly acidic, arguing against a simple recent tandem duplication. Instead, their organization is more consistent with complementary roles within a multicomponent distal-tail apparatus. Complex receptor-binding assemblies containing structurally distinct adaptor, fiber, spike, or distal recognition proteins are well documented among tailed phages.^17^

We therefore propose that WCF59042.1 may function as a proximal adaptor or connector that positions the receptor-interaction machinery at the tail tip, whereas WCF59043.1 may contribute a more distal recognition component. The exceptionally strong C3 prediction for WCF59043.1 is particularly notable because trimeric organization is a recurrent structural feature of phage receptor-binding proteins. However, the present data do not establish the physical orientation of either protein or prove that they directly interact. The proposed tandem organization should therefore be regarded as a structural working model to guide experimental investigation.

### Type IV pilus recognition provides a biologically plausible receptor model

A particularly important observation comes from recent studies of other *P. aeruginosa* phages. Martins et al. demonstrated that phages ZC01 and ZC03 require the type IV pilus for infection of *P. aeruginosa* PA14 and identified PilA as the receptor component recognized by these phages.^10^ Loss of *pilA* markedly reduced or abolished susceptibility, while genetic complementation restored the phage-sensitive phenotype, providing independent evidence that type IV pili are important receptors for diverse *P. aeruginosa* phages.

This finding strengthens the biological plausibility of our UF_RH1 model, particularly when considered together with the previously reported type IV pilus dependence of the closely related DLP phages. Nevertheless, conserved gene organization and receptor usage by related phages do not establish that UF_RH1 necessarily recognizes the same receptor. Receptor-binding proteins are major determinants of host specificity, and comparatively small structural or sequence changes can alter bacterial recognition and host range.^17^ Accordingly, we interpret WCF59042.1 and WCF59043.1 as a candidate type IV pilus-interacting module, while direct UF_RH1 binding to PilA or assembled pili remains to be demonstrated experimentally.

### Structural and mechanistic interpretation

The proposed adsorption model places WCF59042.1 and WCF59043.1 at the interface between the phage tail and the bacterial receptor. Initial recognition may involve engagement of an exposed type IV pilus by the distal receptor-binding complex, followed by stabilization of the virion and movement toward the bacterial surface. Similar principles have been demonstrated structurally in other tailed phages, in which receptor engagement at the distal tail induces conformational rearrangements that promote irreversible attachment and genome delivery.^16^ In bacteriophage lambda, for example, cryo-electron microscopy demonstrated major structural transitions in the trimeric tail-tip receptor-binding machinery after interaction with LamB.^16^

Figures 4 and 5 therefore represent hypothesis-generating models rather than experimentally resolved UF_RH1 protein complexes. They integrate the computational findings into a sequential infection pathway consisting of receptor encounter, adsorption, movement toward the bacterial envelope, genome delivery, intracellular replication, virion assembly, and eventual lysis.

### Transmission electron microscopy supports UF_RH1 morphology and productive infection

Transmission electron microscopy provides complementary ultrastructural support for the biological interpretation of UF_RH1. Purified particles displayed a recognizable capsid associated with a tail, consistent with the morphology expected for a tailed double-stranded DNA bacteriophage. Following infection of *P. aeruginosa*, electron microscopy demonstrated phage particles associated with bacterial cells and accumulation of newly formed particles during infection. Later-stage images showed disruption of the bacterial envelope, release of intracellular material, and extracellular progeny virions, findings consistent with productive replication followed by host-cell lysis.

These observations are relevant to the computational findings because they place the predicted receptor-binding proteins within the context of a morphologically identifiable tailed virion. The WCF59042.1/WCF59043.1 complex is therefore proposed to function during the earliest stage of this experimentally observed infection cycle, whereas downstream genome replication, assembly, and lytic release are mediated by the broader UF_RH1 genomic program.

### Implications for phage biology and engineering

Identification of the molecular determinants of adsorption has implications beyond annotation. Receptor-binding proteins are major determinants of phage host range, and alteration or exchange of RBP domains can modify bacterial specificity. Consequently, structural definition of the UF_RH1 distal-tail apparatus may eventually provide a framework for understanding why closely related phages differ in host range and for rational modification of receptor specificity.

The present results do not demonstrate that WCF59042.1 or WCF59043.1 can yet be used for therapeutic engineering. Rather, they reduce a large set of hypothetical proteins to two experimentally tractable candidates and generate specific hypotheses regarding protein organization, oligomerization, and receptor recognition. Recombinant binding studies, *pilA*-deficient bacterial models, direct PilA or pilus interaction assays, and structural characterization of the WCF59042.1/WCF59043.1 complex will be required to determine whether the proposed model accurately represents the UF_RH1 adsorption machinery.

## Limitations

This study is computational. No biochemical or genetic experiment demonstrates direct binding of WCF59042.1 or WCF59043.1 to P. aeruginosa, PilA, or assembled type IV pili. RBP classifiers can generate false positives, especially at moderate confidence. Seq2Symm predicts symmetry rather than biological function. The available ESMFold/ESMFold2 models are predictions rather than experimentally determined structures. The proposed WCF59042.1/WCF59043.1 complex has not been directly docked or solved. Finally, receptor usage cannot be inferred solely from synteny. These limitations define a clear experimental validation pathway rather than diminishing the value of the computational prioritization.

## Experimental validation priorities

The most direct next experiments as shown in table 2 are: (i) recombinant expression of WCF59042.1 and WCF59043.1; (ii) binding of each protein to susceptible P. aeruginosa cells; (iii) comparison of binding and UF_RH1 adsorption in wild-type, ΔpilA, and complemented strains; (iv) direct PilA or purified pilus interaction measurements by biolayer interferometry or surface plasmon resonance; (v) testing whether soluble WCF59042.1 or WCF59043.1 competitively inhibits phage adsorption; and (vi) biochemical assessment of WCF59043.1 trimerization by SEC-MALS, native mass spectrometry, or cryo-EM.

**Table 1.** Computational evidence supporting reannotation of WCF59042.1 and WCF59043.1.

| **Feature** | **WCF59042.1** | **WCF59043.1** |
| --- | --- | --- |
| **Current GenBank annotation** | Hypothetical protein | Hypothetical protein |
| **Length** | 317 aa | 320 aa |
| **Molecular mass** | 33.50 kDa | 34.99 kDa |
| **Predicted pI** | 8.19 | 5.24 |
| **GRAVY** | -0.073 | -0.033 |
| **RBP prediction** | Tail spike | Tail spike |
| **Predictor** | DepoScope | RBPdetect2 |
| **RBP probability** | 1.00 | 0.51 |
| **Structure resource** | ESMFold | ESMFold2 |
| **Reported pLDDT** | Not used | 73.2 |
| **Predicted oligomeric symmetry** | Not used | C3 |
| **C3 probability** | Not used | 0.990 |
| **DLP2 positional counterpart** | gp22 | gp23 |
| **DLP2 counterpart size** | 320 aa | 321 aa |
| **Historical DLP2 annotation** | Hypothetical/virion protein | Hypothetical/virion protein |
| **Proposed UF_RH1 role** | Proximal tail-spike/connector candidate | Distal trimeric receptor-binding candidate |

**Table 2.** Testable predictions generated by the proposed model.

| **Prediction** | **Experimental test** | **Evidence supporting model** |
| --- | --- | --- |
| **WCF59042.1 associates with the distal tail** | Virion proteomics, immunogold EM, genetic tagging | Synteny and high-confidence TSP prediction |
| **WCF59043.1 forms a trimer** | SEC-MALS, native MS, cryo-EM | Seq2Symm C3 = 0.990 |
| **One or both proteins bind type IV pili** | WT versus ΔpilA binding; purified PilA/pilus binding | DLP1/DLP2 receptor biology |
| **Soluble candidate protein competes with UF_RH1 adsorption** | Adsorption and plaque-inhibition assays | Predicted receptor occupancy |
| **WCF59042.1 and WCF59043.1 form a complex** | Pull-down, BLI/SPR, cross-linking MS | Conserved tandem gene organization |
| **Distal-tail module contributes to host range** | Targeted mutagenesis/domain exchange | RBP biology and sequence divergence |

## Conclusions

Structure-informed reanalysis of Pseudomonas phage UF_RH1 identifies WCF59042.1 and WCF59043.1 as the strongest candidates for a previously unrecognized tandem distal-tail receptor-binding module. WCF59042.1 combines a high-confidence tail-spike classification with a strongly supported C-terminal tail-spike-associated region, whereas WCF59043.1 combines an independent RBP prediction with a strong C3 symmetry signal. Conserved synteny places both proteins in the same positions as unresolved DLP2 virion proteins immediately downstream of the tape-measure gene. Together, these observations support a model in which WCF59042.1 acts proximally as a connector/adaptor and WCF59043.1 contributes a distal trimeric recognition element. The close relationship to type-IV-pilus-dependent DLP1/DLP2 makes pilus recognition an experimentally compelling, but still unproven, hypothesis for UF_RH1.

## Data availability

The complete UF_RH1 genome and associated protein sequences are publicly available under GenBank accession OQ259603.1. Raw sequencing reads from the original genome study are available through BioProject PRJNA936202, BioSample SAMN33343989, and SRA accession SRR23558469. DLP2 is available under GenBank accession KR537871. All sequence-based analyses described here use publicly accessible data.

## Code and supplementary data availability

Physicochemical sequence calculations are reproducible using Biopython version 1.86 and Bio.SeqUtils. ProtParam. Supplementary Table S1 contains the exact input sequences, accessions, coordinates, physicochemical outputs, PhaRBP/RBP prediction outputs, analysis dates, and source links. The accompanying supplementary package includes FASTA inputs, CSV/XLSX outputs, a pinned requirements file, and a Python script that reproduces the ProtParam calculations.

## Author contributions

G.E.: investigation, original phage laboratory work, data review, and manuscript review. M.M.: conceptualization, computational investigation, interpretation, manuscript preparation, and supervision. Contributions should be confirmed by all authors before submission.

## Generative AI statement

Generative artificial intelligence (AI) tools were used to assist with language editing and the development of selected schematic illustrations in this manuscript. AI-generated content was reviewed and verified by the authors for scientific accuracy. AI was not used to generate, modify, or analyze primary experimental data, and all scientific interpretations and conclusions were made by the authors.

## Figure preparation statement

Selected schematic figures and graphical elements were created with the assistance of generative AI and subsequently assembled, edited, labeled, and refined using Adobe Photoshop. AI-generated components were used for illustrative purposes only and do not represent experimentally determined molecular structures unless explicitly stated. Original experimental images, including transmission electron microscopy data, were retained as primary data; image processing was limited to presentation-related adjustments and did not alter the underlying scientific information.

