## Supplementary material for "Comparative genomics and structure-informed analysis identify a putative tandem receptor-binding module in *Pseudomonas* phage UF_RH1": CODE_AND_DATA_AVAILABILITY.txt

CODE AND SUPPLEMENTARY DATA AVAILABILITY

Physicochemical sequence calculations reported in this study can be reproduced using
Python and Biopython version 1.86 (Bio.SeqUtils.ProtParam). The accompanying Python
script, reproduce_physicochemical_analysis.py, uses the exact protein sequences supplied
in Supplementary_Sequences_S1.fasta and generates ProtParam_reproduced.csv.

Supplementary_Table_S1 is provided in both CSV and XLSX formats and contains the protein
accessions, sequence coordinates, physicochemical properties, receptor-binding protein
prediction outputs, and other annotations used in the study.

The receptor-binding and structural predictions reported in the manuscript were retrieved
from publicly accessible computational resources as described in the Methods section.
The deposited files reproduce the sequence-based physicochemical calculations but do not
re-run external web-based prediction services.

Files included:
- Supplementary_Sequences_S1.fasta
- Supplementary_Table_S1.csv
- Supplementary_Table_S1.xlsx
- reproduce_physicochemical_analysis.py
- ProtParam_reproduced.csv
- requirements.txt
- README.md
- MANIFEST_SHA256.txt
