## Supplementary material for "Comparative genomics and structure-informed analysis identify a putative tandem receptor-binding module in *Pseudomonas* phage UF_RH1": README.md

# UF_RH1 code and supplementary data

This package accompanies the manuscript on structure-informed computational reannotation
of *Pseudomonas* phage UF_RH1.

## Reproducing the physicochemical analysis

1. Install Python 3.10 or later.
2. Install the pinned dependency:

   `pip install -r requirements.txt`

3. Run:

   `python reproduce_physicochemical_analysis.py Supplementary_Sequences_S1.fasta`

The script writes `ProtParam_reproduced.csv`.

## Contents

- `Supplementary_Sequences_S1.fasta` — exact amino-acid sequences used for analysis.
- `Supplementary_Table_S1.csv` — supplementary data table in CSV format.
- `Supplementary_Table_S1.xlsx` — supplementary data table in Excel format.
- `reproduce_physicochemical_analysis.py` — reproducibility script.
- `ProtParam_reproduced.csv` — reproduced physicochemical outputs.
- `requirements.txt` — pinned Python dependency.
- `CODE_AND_DATA_AVAILABILITY.txt` — manuscript-ready availability statement.
- `MANIFEST_SHA256.txt` — checksums for integrity verification.

## Scope

The Python code reproduces the sequence-derived physicochemical calculations. External
RBP, structure, and oligomerization predictions reported in the manuscript were retrieved
from the public resources described in the Methods and are recorded in the supplementary
table rather than re-run by this script.
