## Supplementary material for "Comparative genomics and structure-informed analysis identify a putative tandem receptor-binding module in *Pseudomonas* phage UF_RH1": reproduce_physicochemical_analysis.py

```
#!/usr/bin/env python3
"""
Reproduce physicochemical properties reported in Supplementary Table S1.

Requirements:
    Python 3.10+
    biopython==1.86

Usage:
    python reproduce_physicochemical_analysis.py Supplementary_Sequences_S1.fasta

Outputs:
    ProtParam_reproduced.csv
"""
import csv
import sys
from Bio import SeqIO
from Bio.SeqUtils.ProtParam import ProteinAnalysis

if len(sys.argv) != 2:
    raise SystemExit("Usage: python reproduce_physicochemical_analysis.py Supplementary_Sequences_S1.fasta")

input_fasta = sys.argv[1]
out_csv = "ProtParam_reproduced.csv"

headers = [
    "Protein_accession","Length_aa","Molecular_weight_Da","Predicted_pI",
    "Aromaticity","GRAVY","Instability_index","Acidic_D_plus_E_count",
    "Basic_K_plus_R_plus_H_count"
]

rows = []
for record in SeqIO.parse(input_fasta, "fasta"):
    seq = str(record.seq).replace("*","").upper()
    a = ProteinAnalysis(seq)
    rows.append([
        record.id,
        len(seq),
        a.molecular_weight(),
        a.isoelectric_point(),
        a.aromaticity(),
        a.gravy(),
        a.instability_index(),
        seq.count("D") + seq.count("E"),
        seq.count("K") + seq.count("R") + seq.count("H"),
    ])

with open(out_csv, "w", newline="") as f:
    w = csv.writer(f)
    w.writerow(headers)
    w.writerows(rows)

print(f"Wrote {out_csv}")
```
